# CPExtract, a Software for the Automated Tracer-Based Pathway Specific Screening of Secondary Metabolites in LC-HRMS Data

**DOI:** 10.1101/2021.10.20.465085

**Authors:** Bernhard Seidl, Rainer Schuhmacher, Christoph Bueschl

## Abstract

The use of stable isotopically labeled tracers is a long-proven way of specifically detecting and tracking derived metabolites through a metabolic network of interest. While recently developed stable isotope assisted methods and associated, supporting data analysis tools have greatly improved untargeted metabolomics approaches, no software tool is currently available that allows to automatically search LC-HRMS chromatograms for completely free user-definable isotopolog patterns expected for the metabolism of labeled tracer substances.

Here we present Custom Pattern Extract (CPExtract), a versatile software tool that allows for the first time the high-throughput search for user-defined isotopolog patterns in LC-HRMS data. The patterns can be specified via a set of rules including the presence or absence of certain isotopologs, their relative intensity ratios as well as chromatographic co-elution. Each isotopolog pattern satisfying the respective rules is verified on a MS-scan level and also in the chromatographic domain. The CPExtract algorithm allows the use of both labeled tracer compounds in non-labeled biological samples as well as a reversed tracer approach, employing non-labeled tracer compounds along with globally labeled biological samples.

In a proof of concept study we searched for metabolites specifically arising from the malonate pathway of the filamentous fungi *Fusarium graminearum* and *Trichoderma reesei*. 1,2,3-^13^C_3_-malonic acid diethyl ester and native malonic acid monomethyl ester were used as tracers. We were able to reliably detect expected fatty acids and known polyketides. In addition, up to 189 and 270 further, unknown metabolites presumably including novel polyketides were detected in the *F. graminearum* and *T. reesei* culture samples respectively, all of which exhibited the user-predicted isotopolog patterns originating from the malonate tracer incorporation.

The software can be used for every conceivable tracer approach. Furthermore, the rule sets can be easily adapted or extended if necessary. CPExtract is available free of charge for non-commercial use at https://metabolomics-ifa.boku.ac.at/CPExtract.

## INTRODUCTION

Untargeted metabolomics aims at the unbiased detection of all metabolites produced by a biological system that are covered by the respective analytical method used. Typically both qualitative and quantitative differences in metabolites resulting from natural fluctuations, genetic modifications (e.g. WT versus knock-out) or experimental biotic or abiotic stress factors (e.g. untreated versus treated in any form) are then examined [1]. Nowadays, besides nuclear magnetic resonance (NMR) spectroscopy, liquid chromatography coupled with high-resolution mass spectrometry (LC-HRMS) is the most commonly used analytical technique to cope with this task. LC-HRMS allows the detection of a large number of different metabolites in the samples. However, usually only few of these can be identified by comparison with reference standards [2]. Moreover, it is often not even possible to reliably differentiate between signals deriving from contaminants or artefacts and those of truly biological metabolites [3]. To encounter this problem, stable isotopically labeled samples, biological material or tracer molecules can be employed. The ingenious basic principle here is that labeling is incognito for the organism, i.e. labeled substances are metabolized almost the same way as unlabeled substances in biochemical reactions, but the label can be easily traced in analytical measurements [4, 5]. Stable isotope-assisted (SIA) workflows were used as early as the 1950s in comprehensive studies of metabolic pathways, where the progression of biochemical signaling pathways and metabolic intermediates was investigated by tracking changes in the isotopic composition of metabolites of the metabolic pathway under study over time. This made it possible to study metabolic pathways and networks or to investigate the metabolic fate of substances (e.g. drugs or toxins) into intermediate or end products. Typically, _13_C, _15_N or _2_H isotopes are used for tracking derived metabolites through a given metabolic network of interest [5-9].

Additionally, the labeling information can also be used for improved unknown compound annotation, e.g. by means of interpretation of SIA tandem mass spectrometry data [10].

Meanwhile software-supported approaches for the untargeted search for tracer-derived compounds also are commonly used in metabolomics, where in contrast to the more specific biosynthetic pathway studies, a snapshot of the entire metabolic state that can be captured by the method used is analyzed [1]. In the latter case, the artificial isotope patterns caused by the metabolism of the labeled tracer are used to reliably track all tracer-derived metabolites that contain the entire tracer or parts thereof.

However, especially in untargeted SIA metabolomics studies, the huge amounts of raw LC-HRMS data do not allow the comprehensive, manual evaluation anymore. Therefore, a variety of (semi)automated SIA methods and data evaluation tools have recently been developed and greatly helped to improve untargeted metabolomics approaches. Examples of such tools are MetExtract II [11], X13CMS [12], HiTIME [13], geoRge [14], ALLocator [15], NTFD [16], or CompoundDiscoverer 3.x (Thermo Scientific). Each of these tools has been developed for particular isotope patterns or is based on direct comparison of native and labeling-derived isotope signatures and is therefore limited to its respective specific application. For example, MetExtract II [11] and HiTIME [13] both require LC-HRMS data sets consisting of native and uniformly _13_C-labeled compounds and thus detect either such labeled metabolites or their bio-transformation products. On the other hand, X13CMS [12], geoRge [14] or NTFD [16] detect compounds based on deviations from native or natural isotopolog patterns after treatment of biological systems with an isotopically labeled precursor (such as U-^13^C_6_ glucose) in comparison to the same samples treated with the non-labeled precursor. To the best of the authors’ knowledge however, there is no software tool available to date, that allows the user to freely and flexibly specify a custom isotopolog pattern, which is then searched for in the LC-HRMS data set. Here we present Custom Pattern Extract (CPExtract) that allows the user to specify custom isotopolog patterns together with pre-set relative intensity ranges to be searched for, thereby enabling to specifically filtering for metabolite ions of interest that agree with the desired and diagnostic isotopolog pattern.

## METHODS

### 1. CPExtract application example

The application of CPExtract is presented by searching specifically for metabolites of the malonate pathway, namely fatty acids, polyketides, and possible hybrid metabolites thereof (e.g. nonribosomal peptide-polyketides or prenylated polyketides) in culture samples of the filamentous fungi *F. graminearum* and *T. reesei*.

Two complementary tracer approaches were used. First 1,2,3-^13^C_3_-malonic acid diethyl ester was used as tracer substance. This approach will henceforth be referred to as the standard tracer approach, since the tracer was isotopically labeled (_13_C) and the cultivation was carried out with native glucose as the sole carbon source in the growth medium.

The second tracer substance was malonic acid monomethyl ester. Since this substance is not readily available in a _13_C isotopically labeled form, a complementary approach was chosen, which will henceforth be referred to as the reversed tracer approach. Here the native form of malonic acid monomethyl ester was used as a tracer, while the only other available carbon source for the fungus was U-^13^C_6_-labeled glucose. Detailed information about the cultivation workflow can be found in the supplementary information.

### 2. CPExtract algorithm

The automated CPExtract workflow is based on the MetExtract II software, which originally has been designed to detect pairs of native and uniformly _13_C-labeled metabolite ions. CPExtract replaces this fixed and rigid definition by a set of freely definable rules by which the isotope patterns to be searched for can be arbitrarily defined by the user. The individual rule set is defined via a Python vector of objects where each is an instance of one of the currently available “Rule” objects, which include presence or absence of certain isotopologs as well as abundance or area-ratios between different isotopologs. Further custom user-defined rules can be easily added by defining additional subclasses. Automatic data processing then filters for all signals with isotope patterns that correspond to the defined set of rules. A schematic representation of the data processing workflow and algorithm is shown in Figure 1.

**Figure 1.**
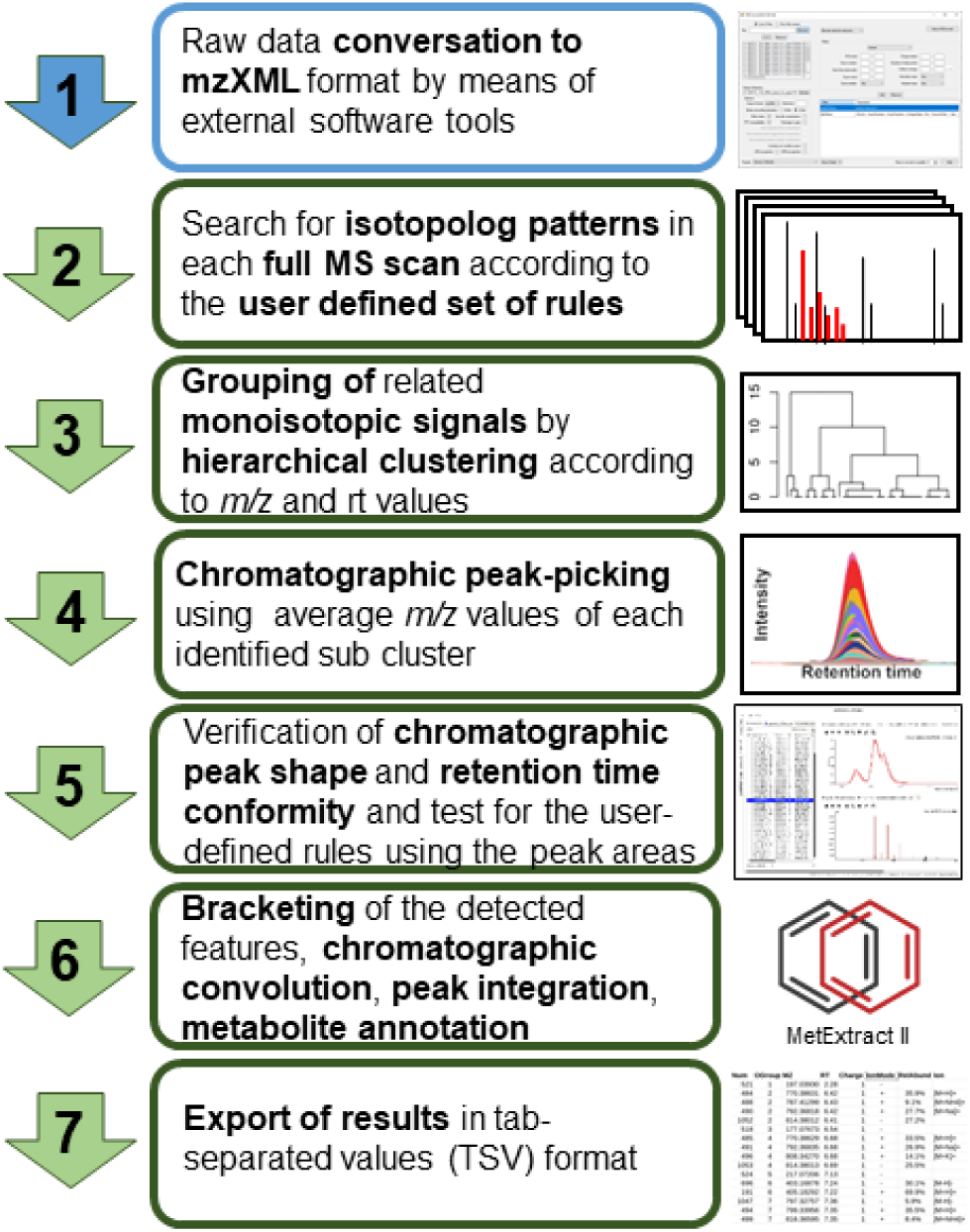
Schematic overview chart of the data processing workflow and CPExtract algorithm.

The individual steps of the processing workflow and CPExtract algorithms are:

1. In order to be processed the manufacturer-specific raw data chromatograms must be converted into the mzXML format with e.g. ProteoWizard MSConvert [17].
2. Data processing in CPExtract starts at the scan level, whereby each individual *m/z* signal in a full scan mass spectrum is initially considered to represent the principal isotopolog of a putative metabolite ion of interest (henceforth termed X). This assumption is then verified by checking the user defined set of rules. Currently, the following distinct rules are available: The “PresenceRule” (a particular isotopolog must be present and be within expected ratios relative to other isotopologs) and “AbsenceRule” (a specified isotopolog must not be present). For these two rules a “minIntensity” parameter, which can be used to define a minimum signal intensity that the isotopolog must have and a “verifyChromPeaksSimilarity” parameter, which specifies that the chromatographic peaks of the isotopologs must be present and co-elute. Furthermore the “RatioRule” (checks whether an intensity ratio of two isotopologs or even two groups of several summed up isotopolog intensities corresponds to a certain defined ratio or checks if both ratios are within a defined range), the “AnyIntensityRule” and the “AllIntensityRule” (either any or all specified isotopologs must exceed a certain intensity threshold) are available. Moreover, users can easily implement their own rules if required. It is then checked if the isotopologs defined in the rule set relative to the assumed X isotopolog are valid (e.g. if certain isotopologs are present or absent). Only if X and all its putatively related isotopolog peaks meet the specified rules, X is used for subsequent data processing.
3. All MS signals (i.e. Xs) that fulfilled the rules in the previous step are then clustered using hierarchical clustering. Sub-clusters of the thereby generated dendrogram within a low mass deviation window (scan-to-scan variability) are kept. This dendrogram cutting step also takes the acquisition time (scan number according to the retention time) of the MS-scans into consideration so that isomers end up in different sub-clusters.
4. Data processing then continues on the chromatographic level. Therefore, an average *m/z* value is calculated for Xs of each sub-cluster present from the previous step, and peak picking is carried out using the continuous wavelet transform (CWT)-based centWave algorithm [18].
5. Only those isotope pattern candidates (or Xs) identified in the previous steps, whose chromatographic retention and peak shape show sufficient agreement with the user’s rules, are ultimately accepted as hits. For this, all chromatographic peaks of the isotopologs must show perfect chromatographic co-elution, which is tested via the Pearson correlation coefficient, and additionally also the isotopolog ratios based on the chromatographic peak areas are again checked and must meet the user’s rules-criteria.
6. The remaining steps of CPExtract are the same as implemented in MetExtract II [11], namely bracketing of the detected features across all samples, chromatographic convolution (i.e. grouping and annotation of metabolite ions from the same metabolite), and peak area re-integration.
7. The results of the automated data processing are provided in the form of a tab-separated values (TSV) file containing a data matrix in which all detected principal isotopologs (Xs). The results can also be visualized and checked in the software’s graphical user interface, whereby both the extracted ion chromatograms and the spectra are illustrated.

### 3. Fungal strains and cultivation conditions

*Fusarium graminearum* PH-I (wild-type) obtained from Gerhard Adam, BOKU, Institute for Microbial Genetics (IMiG) and three different *Trichoderma reesei* strains, namely QM6aΔ*tmus53* (wild-type) [19], QM6a Δ*tmus53*Δ*sor1* (a knock-out mutant lacking the gene encoding the SOR1 polyketide synthase, the first enzyme in the sorbicillinoid biosynthetic pathway) [20] and QM6a Δ*tmus53ReYpr1* (a yellow pigment regulator 1 transcription factor over expressing strain) [21] were used for the preparation of culture medium supernatant and mycelium samples. To obtain the data sets, the samples were measured with LC-HRMS. Detailed information about the SIA-tracer workflow, cultivation, sample preparation and measurement are given in the Supporting information.

### 4. Conversion of LC-HRMS raw data files

The raw data files were converted into mzXML format using ProteoWizzard MSConvert Software [17] version 3.0.19210 (settings: mzXML output format, 32-bit binary encoding precision, enabled index writing, enabled TPP compatibility, disabled zlib and gzip compression, peak picking using vendor algorithm).

## 5. CPExtract rules and parameter settings for data processing

### 5.1. Processing of LC-HRMS data generated with the standard tracer approach

The following set of rules was used to detect isotopolog patterns of fully ^12^C metabolites that incorporate several ^13^C_2_ units derived from the _13_C-labeled malonic acid diethyl ester tracer as expected from the standard tracer approach:

X represents the native, monoisotopic form of the metabolite ions and consists only of _12_C isotopes for each carbon atom and X_+υ_ indicates that υ ^12^C atoms are exchanged for _13_C atoms in that particular isotopolog relative to X.

- X must have a signal abundance of at least 1E5 counts. X_+2_ and X_+4_ must have a signal intensity of at least 5E4 counts. (part of PresenceRule)
- As X already represents the fully _12_C isotope form of the metabolite, the signals for putative ^12^C-_1_ and ^12^C-_2_ isotopologs should therefore not be present. To avoid false negative results (e.g. signal noise), signals with an intensity value of maximum 5 % of X are accepted for both isotopologs without a rule violation. (AbsenceRule)
- The metabolite ion containing one and two _13_C-malonate tracer derived extension units X_+2_ and X_+4_ must be present and their abundance has to be within the range of 10 to 300 % of X and X_+2_ respectively. (RatioRule)
- Moreover the metabolite ions containing one or two partial isotopically labeled extension units (respectively one ^13^C and one ^12^C) X_+1_ and X_+3_ must be present and must have an abundance of 10 to 200 % relative to X and X_+2_as well as X_+2_ and X_+4_ respectively. (RatioRule)
- The ratio of the abundance of X totaled with the abundance of X_+2_ to the abundance of X_+1_ must be the same as of the abundance ratio of X_+2_ totaled with the abundance of X_+4_ to the abundance of X_+3_ with a maximum permissible deviation of these ratios of ± 10 % respectively (i.e. (X+X_+2_)/X_+1_ ≈ (X_+2_+X_+4_)/X_+3_). (RatioRule)
- The isotopologs X, X_+1_, X_+2_, X_+3_ and X_+4_ must be present as co-eluting chromatographic peaks (Pearson correlation ≥ 0.85). (part of first PresenceRule)

### 5.2. Processing of LC-HRMS data generated with the reversed tracer approach

To detect isotopolog patterns of fully _13_C labeled metabolites that incorporate one or more ^12^C_2_ units derived from the native malonic acid monomethyl ester, slightly modified rules compared to the standard tracer approach were used.

In case of the reverse tracer approach, X represents the uniformly _13_C-labeled form of the metabolite ions and consists only of ^13^C isotopes for each carbon atom and X_-υ_ indicates that υ _13_C atoms are exchanged for _12_C atoms.

- X must have a signal abundance of at least 1E5 counts. X_-2_, X_-4_ and X_-6_ must have a signal intensity of at least 5E4 counts. (PresenceRule)
- Since here X already represents the fully labeled _13_C isotope form of the metabolite, the signals for putative ^13^C_+1_ and ^13^C_+2_ isotopologs should not be present. Signals up to 5 % of X were tolerated. (AbsenceRule)
- The metabolite ion containing one, two and three ^12^C-malonate tracer derived extension units X_-2_, X_-4_ and X_-6_ must be present and their abundance has to be in a range of 10 to 300 % of X, X_-2_ and X_-4_ respectively. (RatioRule)

Moreover, the metabolite ions containing one or two extension units with one ^12^C and one ^13^C (i.e. X_-1_, X_-3_ and X_-5_) must be present and must have an abundance of 10 to 200 % relative to X and X_-2_, X_-2_ and X_-4_, and X_-4_ and X_-6_ respectively. (RatioRule)

- The ratio of the abundance of X totaled with the abundance of X_-2_ to the abundance of X_-1_ must be the same as of the abundance ratio of X_-2_ totaled with the abundance of X_-4_ to the abundance of X_-3_ as well as the abundance ratio of X_-4_ totaled with the abundance of X_-6_ to the abundance of X_-5_ with a maximum permissible deviation of these ratios of ± 10 % respectively (i.e. (X+X_-2_)/X_-1_ ≈(X_-2_+X_-4_)/X_-3_ ≈ (X_-4_+X_-6_)/X_-5_). (RatioRule)
- All isotopologs defined via “PresenceRule” instructions (X, X_-1_, X_-2_, X_-3_, X_-4_, X_-5_ and X_-6_) had to be present as coeluting chromatographic peaks and must show a highly similar peak shape (Pearson correlation ≥ 0.85). (part of first PresenceRule)

### 5.3. General processing parameters

The further data processing parameter settings of CPExtract were: intra-scan-mass-accuracy: ±5 ppm, inter-scan-mass-deviation: ±8 ppm, EIC-extraction window: ±5 ppm, retention-time-window: 3-36 minutes, minimum/maximum chromatographic peak width: 5-25 seconds, minimum Pearson correlation for co-eluting chromatographic peaks: 0.85, maximum allowed retention-time-deviation for bracketing of multiple measurements: 0.1 minutes, maximum allowed *m/z* deviation for bracketing: 5 ppm.

## RESULTS AND DISCUSSION

A novel software tool, named CPExtract, for the comprehensive and automated search for isotope patterns in LC-HRMS data that can be freely defined by the user is presented. Other than existing software tools, it provides the user with an automated framework to mine the LC-HRMS data for certain characteristic isotopolog pattern via a set of predefined and extensible rules. Only isotopolog patterns obeying these rules are reported. Moreover, the large flexibility in defining the target patterns enables to use the CPExtract algorithm in two complementary ways, a standard tracer-, as well as a reversed tracer approach. As a proof of concept, the CPExtract software was used to search specifically for fungal metabolites of the malonate pathway, namely fatty acids and polyketides as well as their derivatives like e.g. nonribosomal peptide-polyketide (NRP-PK) or prenyl-polyketide hybrids in LC-HRMS data.

### 1. Expected theoretical isotope patterns

#### 1.1. Standard tracer approach

Figure 2 shows a typical theoretical isotopolog pattern of a substance which is to be expected for the standard tracer approach, due to the accidental incorporation of both added (^13^C-) tracer-derived and native fungus-produced C_2_ building blocks into its carbon skeleton. Such a pattern can be expected when the bioavailability and incorporation rate of tracer molecules is small in relation to the amount of malonyl-CoA units, endogenously formed from native glucose and used by the fungus itself for biosynthesis.

**Figure 2.**
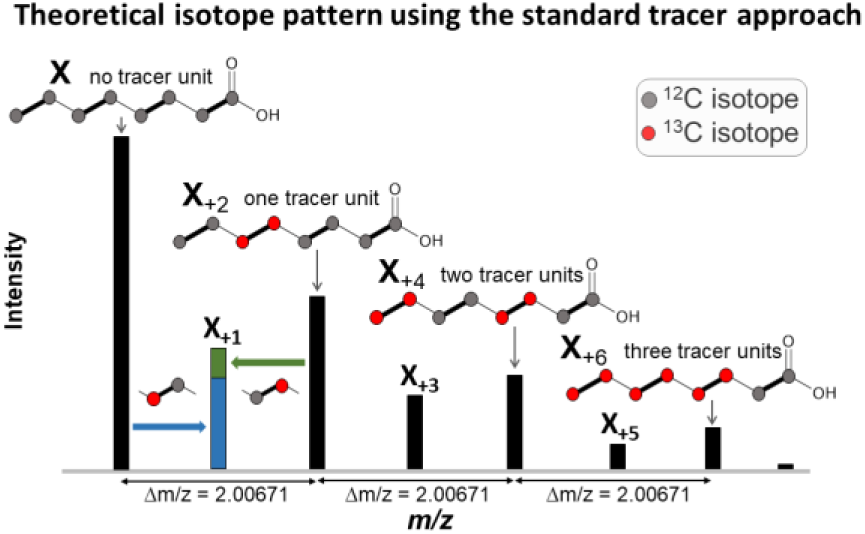
Theoretical isotope pattern of caprylic acid, which can be expected as a result of the incorporation of C_2_-extension units derived from both, native malonate (synthesized by the fungus itself) and stable isotopically-labeled malonate (tracer added to the medium) into the growing hydrocarbon chain.

A staircase shaped isotope pattern is characteristic for continuously produced metabolites, corresponding to the decreasing probability that an increasing number of tracer-derived C_2_ units become incorporated into the synthesized metabolite.

In addition, also the X_+1_, X_+3_ and X_+5_ isotopologs are of high diagnostic value. They result only from the proportion of non-monoisotopic tracer derived extension units incorporated into the metabolites that occur due to the natural abundance of _13_C isotopes in the native carbon source and the _12_C isotopic impurity of the _13_C labeled tracer compound used. Therefore, their signal intensities must be smaller than the mean value of the two neighboring isotopologs with even numbered mass increment (e.g. X_+1_ < (X+X_+2_)/2). Furthermore, the resulting intensity ratio (e.g. (X+X_+2_)/X_+1_) must be almost the same for all analogous isotopolog pairs (e.g. (X_+2_+X_+4_)/X_+3_). The intensity ratio of odd-numbered isotopologs to their even-numbered neighbors is independent of the number of tracer-derived extension units incorporated into the particular metabolite ion, making them ideal for highly specific filtering.

#### 1.2. Reversed tracer approach

Often _13_C-labeled substances are expensive or not available at all. In this case, the reversed approach can be a solution to the problem. When combined with global _13_C labeling of the biological system, the use of a native tracer such as malonic acid monomethyl ester results in a typical isotope pattern that is a mirror image inversion of the pattern expected from the standard tracer approach (Figure 3).

**Figure 3.**
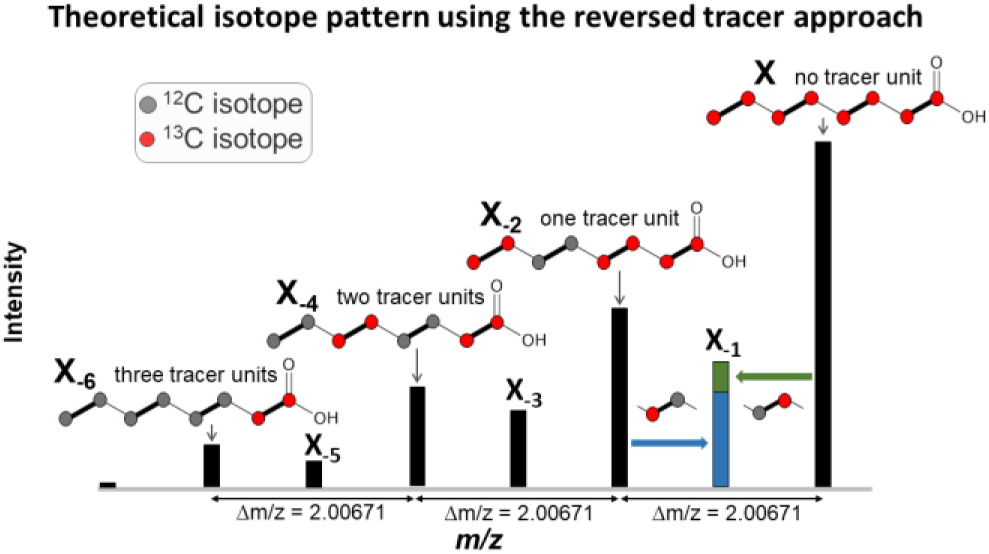
Theoretical isotope pattern of caprylic acid, which can be expected as a result of the incorporation of C_2_ units derived from both, _13_C malonate (synthesized by the fungus itself as grown on U-_13_C_6_-glucose as sole carbon source) and native malonate (tracer added to the medium) into the growing hydrocarbon chain.

Due to the high flexibility in defining rules for the isotope patterns to be searched for, the CPExtract software is able to deal with both tracer approaches equally.

### 2. Exemplification of isotope pattern and tolerance intervals for oleic acid

Figures 4 and 5 show the spectra of oleic acid experimentally measured in *F. graminearum* mycelium extract samples together with the identified monoisotopic isotopolog (X) and the respective tracer-derived isotopologs exemplarily for standard tracer- and reversed tracer approach respectively. For *T. reesei* mycelium extract similar patterns were observed (data not shown). In both illustrations the red boxes show the required isotopologs as well as their permissible intensity ranges as defined in the respective rule set for the standard- and the reversed tracer approach.

**Figure 4.**
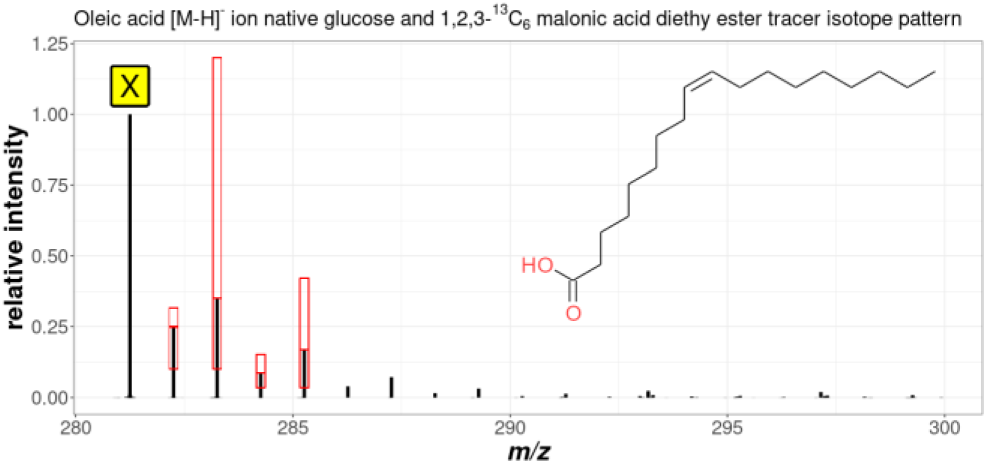
Measured spectrum of the [M-H]^-^ ion of oleic acid in the mycelium extract samples of *F. graminearum* using the standard tracer approach. X denotes the predominant, monoisotopic, native deprotonated molecule ion. Also shown in red boxes are the isotopologs defined in the rules that must be present and their permitted intensity ranges

**Figure 5.**
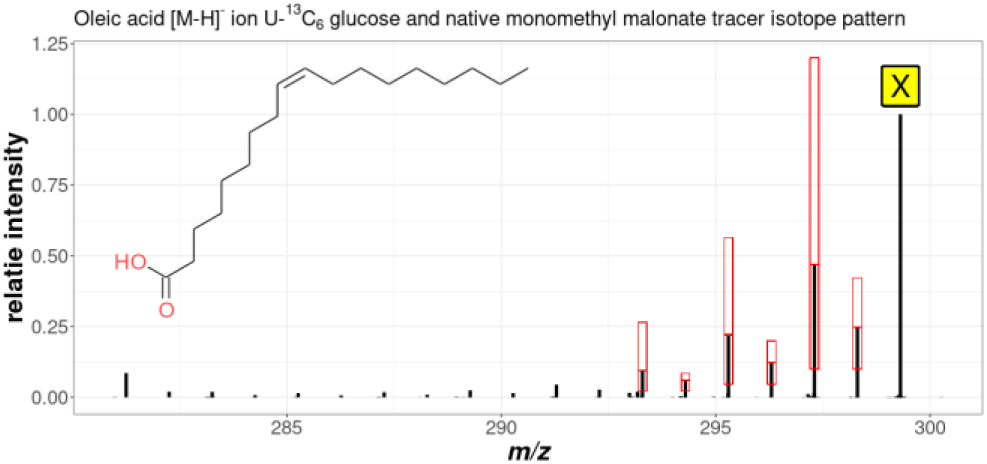
Measured spectrum of the [M-H]^-^ ion of oleic acid in the mycelium extract samples of *F. graminearum* using the reversed tracer approach. X denotes the predominant, monoisotopic fully _13_C labeled deprotonated molecule ion. Also shown in red boxes are the isotopologes defined in the rules that must be present and their permitted intensity ranges

As can be seen from the figures, the rules selected are suitable for the detection of the isotope patterns in the measurement data caused by the random incorporation of tracer-derived C_2_ extension units. Due to the rapidly decreasing signal intensity of isotopologs with an increasing number of tracer derived extension units, not all theoretically existing isotopologs were detected.

For this reason, the rule set of the standard tracer approach required the presence and permissible intensity of at least two C_2_ extension unit isotopologs (X_+2_ and X_+4_) as well as the isotopologs X_+1_ and X_+3_ in between for a hit. In addition, the intensity ratios of the sum of X plus X_+2_ to X_+1_ and X_+2_ plus X_+4_ to X_+3_ respectively had to be equal with a maximum permissible deviation of ±10 %.

With the reversed tracer approach more abundant signals for the X_-n_ isotopolog signals (X_-1_, X_-2,_ …) were obtained compared to the standard tracer approach. Therefore, the rule set was extended by an additional pair of isotopologs corresponding to a third tracer-derived C_2_ extension unit (X _-5_ and X_-6_), further increasing the specificity.

### 3. Isotope patterns found for the polyketides aurofusarin and sorbicillinol

The experimentally observed isotopolog patterns after data processing with CPExtract are exemplified with the two known polyketides aurofusarin for *F. graminearum* and sorbicillinol for *T. reesei*. Results are shown for both tracer approaches in Figure 6.

**Figure 6.**
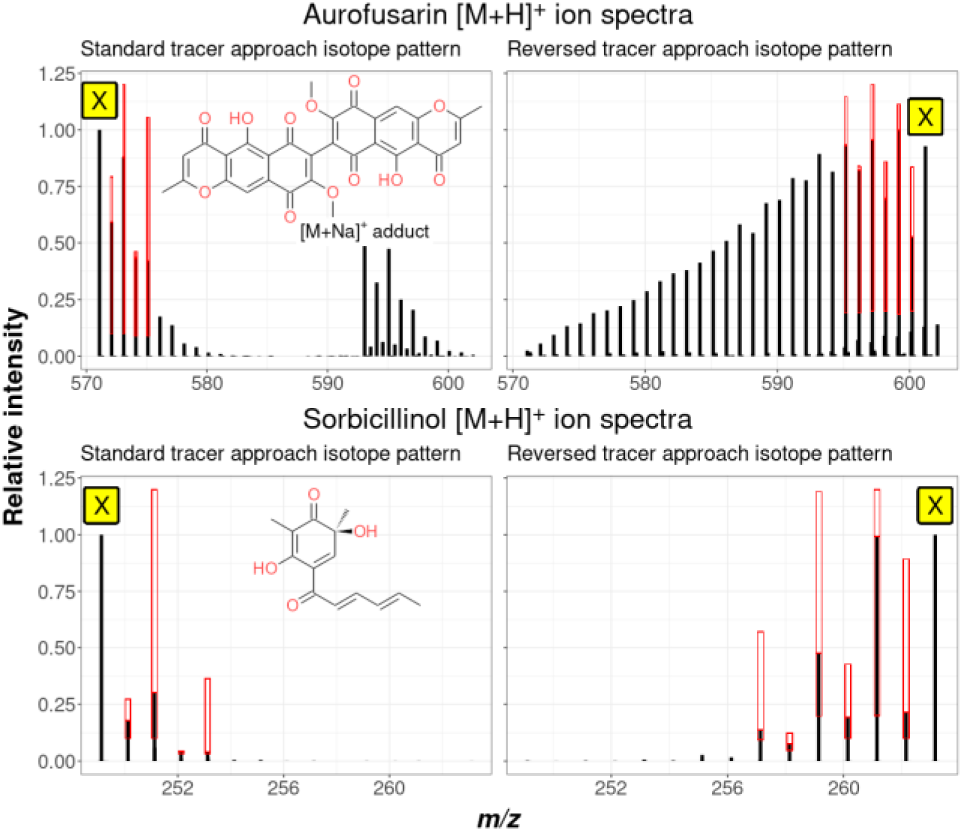
Spectra of the metabolites aurofusarin (*F. graminearum*) and sorbicillinol (*T. reesei*), actually found with CPExtract for both tracer approaches. Also shown in red boxes show the isotopologs defined in the rules that had to be present and their permitted intensity ranges.

The spectrum of aurofusarin from the reversed tracer approach showed all fifteen isotopolog signals corresponding to the incorporation of all fifteen possible malonate tracer-derived C_2_ extension units into the carbon skeleton. In contrast, for aurofusarin in the standard tracer approach and for sorbicillinol in both approaches, only some of the theoretically possible isotopologs were sufficiently abundant for being detected.

### 4. Overall pattern shapes found for the polyketide aurofusarin

In both approaches, the intensity distribution does not necessarily have to follow the strict declining or increasing pattern shown above. Depending on the onset and time period of the metabolite synthesis during cultivation and the prevailing ratio of added tracer and extension units formed by the fungus from the general carbon source, a different intensity profile can also appear. Moreover, the shape of the pattern does also depend on whether a metabolite is possibly composed of two or more building blocks that were initially synthesized independently, as well as the cell compartment localization and time and rate of excretion of the bio synthesized product. Figure 7 illustrates for example the slightly differing isotope patterns obtained for the *F. graminearum* derived polyketide aurofusarin (C_30_H_18_O_12_) in mycelium extract and supernatant samples.

**Figure 7.**
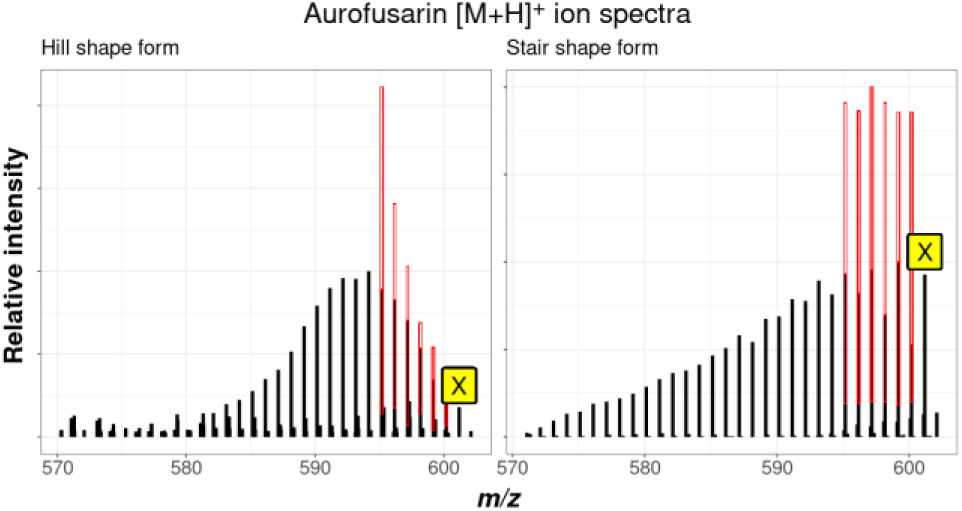
Different isotope pattern shapes of [M+H]_+_ isotopologs of *F. graminearum* derived aurofusarin. A hill shape form was found in mycelium extracts (left) whereas a stair shape form was found in supernatant samples (right).

Consequently, the rules for the permissible intensity ranges for the X_+/-2_ isotopologs must allow adequate leeway for being able to detect all metabolite ions of the target substance class.

While the large tolerance windows needed for the initial pattern search can potentially compromise the filtering specificity and thus lead to false positive search results, additional pattern shape characteristics can be used for further filtering of candidate ions.

Insufficient predictability of the overall shape of the pattern does not affect the respective intensities of the isotopologs located between two adjacent even-numbered isotopologs (e.g. for X_+1_ between X and X_+2_). As already explained above, the odd-numbered (X_±1_, X_±3, …_) isotopologs always result from the sum of the respective isotope fractions and the number of carbon atoms in the metabolite. Furthermore, and perfectly usable as a reliable selection criterion, the relative ratio of the intensity of these isotopologs to that of the respective two analogous neighboring isotopologs with a differing number of tracer derived extension units must remain the same. Therefore, much smaller tolerable intensity intervals can be defined in the rules for the odd-numbered isotopologs without running the risk to miss target metabolite ions. To this end, the initially created candidate list can be refined by checking the constant relative intensities as described above to sort out remaining false positives very reliably.

### 5. Global screening of all tracer-derived metabolite ions

The big advantage of filtering MS features according to certain metabolite classes by CPExtract using specific isotope patterns derived from the corresponding tracer can clearly be seen in Figure 8.

**Figure 8.**
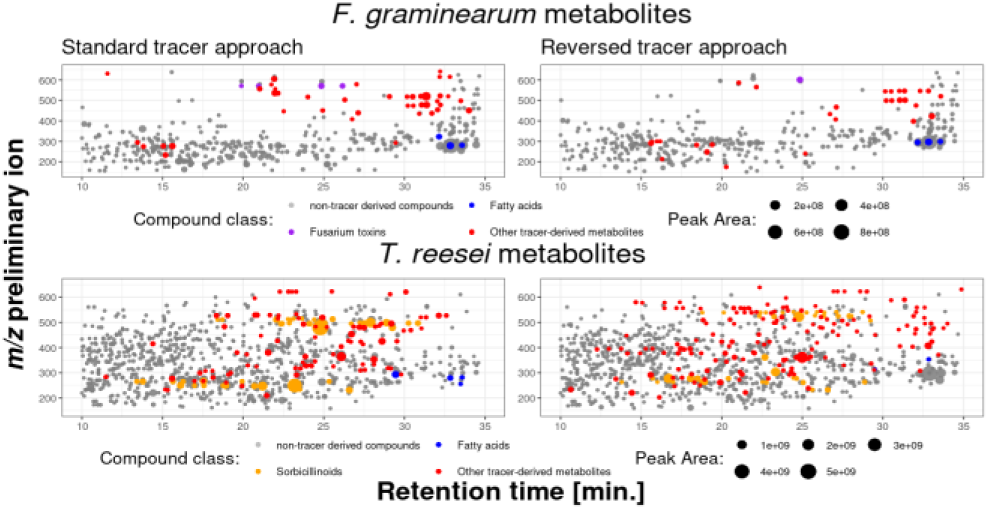
Mass-retention time plot of the dominant ion of each feature group from all true fungal metabolites found by MetExtract II software (shown in gray) in the sample data sets of *F. graminearum* and *T. reesei* (all three genotypes) culture samples. Metabolites detected by CPExtract are shown in purple (known *F. graminearum* metabolites), yellow (known *T. reesei* metabolites), or red (unknown, potential novel tracer-derived metabolites), respectively.

In the *F. graminearum* data set, the MetExtract II evaluation of the global labeling approach resulted in a total number of 621 metabolites found both in supernatant and mycelium extract samples of the wild-type strain PH-I that were actually produced by the fungus (Figure 8). The CPExtract algorithm detected a subset of tracer-derived metabolites from this set, comprising 182 metabolites for the standard tracer approach and 191 metabolites for the reverse tracer approach. This clearly demonstrates the filter effect of the tracer approach combined with the automated CPExtract data evaluation on the targeted metabolic pathway.

In the *T. reesei* data set, consisting of supernatant and mycelium extract samples of each of the three different strains, 1165 metabolites truly originating from the fungus were found by MetExtract II data evaluation. The CPExtract algorithm data processing yielded 167 tracer tracer-derived metabolites for the standard tracer approach and 283 for the reversed tracer approach respectively (Figure 8).

The relatively high number of (malonate derived) metabolites can be explained on the one hand by the combination of three different fungal genotypes and is on the other hand presumably due to the high reactivity of the sorbicillinoids. Members of this class of polyketides are known to spontaneously react non-enzymatically into various other substances or reaction by-products. Figure 9 shows for both tracer approaches all found feature groups detected in the *T. reesei* culture samples. In the Venn diagrams it can be seen that no metabolites annotated as sorbicillinoids were found in the ΔSor1 strain samples, which is as expected since this strain is deficient in the sorbicillinoid polyketide gene cluster and thus not able to produce any sorbicillinoids at all. Furthermore, it can be seen that some additional sorbicillinoids (namely bisorbicillinol, epoxysorbicollinol, sorbiquinol, bisvertinolone, dihydrobisvertinolon) were found exclusively in samples of the overexpressing ReYpr1 strain.

**Figure 9.**
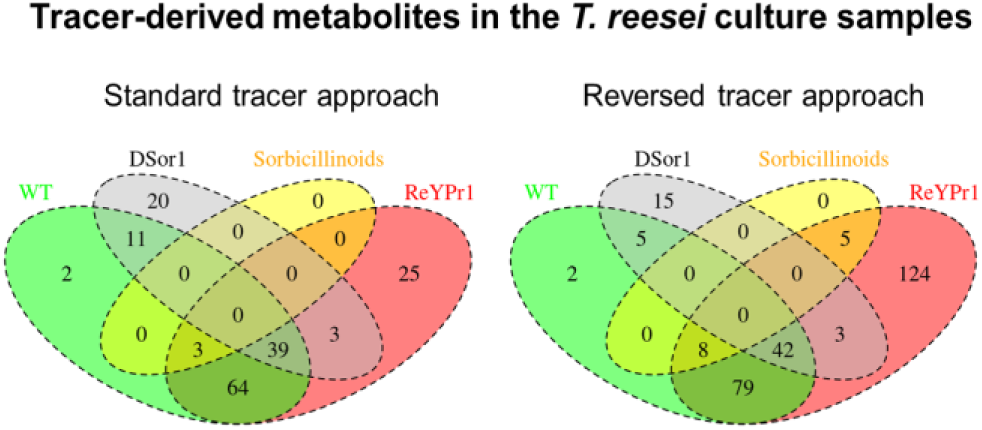
Venn diagrams of all metabolites found with both, the standard tracer and the reversed tracer approach in the investigated *T. reesei* strain culture samples (wildtype (WT) in green, Ypr1-overexpressing ReYpr1 strain in red and Δ*sor1* deletion strain in grey). The annotated sorbicillinoids are shown in yellow.

### 6. Selectivity of CPExtract algorithm

Furthermore, to test the selectivity of the CPExtract algorithm and the rule sets used, culture samples without the addition of the tracer compounds were measured and processed using the same settings. This was done for both the standard- (^12^C glucose) and reversed (^13^C_6_-glucose) approach. In these negative control samples a low number of hits (false-positives) were detected by CPExtract, demonstrating the high specificity of the presented approach. Surprisingly, only in some *T. reesei* negative control samples corresponding tracer-derived isotope patterns were detected. In total, seven corresponding tracer-derived patterns (different ones in each replicate) were detected in supernatant samples of the Ypr1 overexpressing ReYpr1 strain and one tracer-derived pattern was detected in a single replicate of the WT supernatant samples. Manual inspection of the raw data files revealed that the false positive signals were only detected immediately after the measurement of samples, which showed high signal intensities for exactly those ion species. Therefore, it was concluded that the false positives resulted from carryover into successive injections via the insufficiently rinsed injection needle of the HPLC system. Thus, the false positive hits were not caused by a low selectivity of the CPExtract algorithm but correctly detected true polyketides (data not shown).

## CONCLUDING REMARKS

Based on the exemplary search for metabolites of the malonate pathway in the culture samples of two different filamentous fungi using two complementary tracer approaches, it was possible to demonstrate the reliable function of the CPExtract algorithm and its universal and versatile applicability. Through the detection of unknown metabolites, specifically from the substance group of interest, in addition to the known and expected metabolites of the respective fungus, it is evident that the CPExtract software can also benefit the search of various other putatively novel or even completely unknown metabolites. Especially the timely but still demanding topic of natural product discovery is a promising field of application of CPExtract. As exemplified in this article, the reliable filtering the MS data for the fatty acids and polyketides searched for, enables a drastic reduction in the complexity of the global metabolite profiles. This can help to greatly simplify manual data curation that is subsequently required.

CPExtract shows also great potential for other natural product classes such as nonribosomal peptides or terpenoids when being combined with other tracer compounds such as amino acids or mevalonate. Another interesting option is to screen for functionalization of natural products like for example methylations by the use of L-methionine-(methyl- _13_C). Last but not least the metabolic fate of (labeled) potentially toxic xenobiotics can be probed by CPExtract either with the standard or reverse approach.

We anticipate that the presented tracer-approach in combination with the flexible and easy-to-use CPExtract software will be of interest to the metabolomics community working in related fields of research.

CPExtract is available free of charge for non-commercial user at https://metabolomics-ifa.boku.ac.at/CPExtract. Likewise, the LC-HRMS data can be obtained from https://metabolomics-ifa.boku.ac.at/CPExtract/#datasets.

## ASSOCIATED CONTENT

### Supporting Information

The Supporting Information is available free of charge on the ACS Publications website. One section with additional text and three figures describing the SIA-tracer workflow, cultivation, sample preparation and measurement in detail and showing screenshots of CPExtract software (PDF).

## AUTHOR INFORMATION

### Author Contributions

BS and RS designed the experiments. BS carried out the biological experiments and the LC-HRMS analysis. BS and CB implemented the software tool and processed the data set. All authors contributed to the biological interpretation and the implementation of the manuscript.

### Notes

The authors declare no conflict of interest.

## ACKNOWLEDGMENT

The authors would like to thank the Austrian Science Fund (projects ReSMe P-26733, playNICE ZK-74) and the Provincial Government of Lower Austria (project OMICS 4.0) for funding this work.

Furthermore we would also like to thank Robert Mach, Astrid Mach-Aigner, Christian Derntl (TU-Wien) for providing the T.reesei strains and Gerhard Adam and Gerlinde Wiesenberger (University of Natural Resources and Life Sciences, Vienna) for providing the F. graminearum strain.

