## Supporting information for "CPExtract, a Software for the Automated Tracer-Based Pathway Specific Screening of Secondary Metabolites in LC-HRMS Data"

### 1 Stable isotope-assisted (SIA) tracer workflow

#### 1.1 Overview

In the following, the stable isotope-assisted (SIA) tracer workflow briefly is described, which was used for obtaining the culture samples leading to the presented demonstration data sets. The basic scheme of the SIA tracer workflow is shown in Figure S1.

First step was a pre-cultivation of the respective fungus from an inoculum made from fresh fungus spores in minimal liquid medium on a small scale (24 well cell culture plates), initially without adding the corresponding labeled tracer substance. This served to first build up enough biomass without having to worry about a possible influence from the tracer substance.

As soon as the fungus has grown sufficiently (i.e. the fungal culture was in the exponential growth phase), the tracer substance was added. The optimal tracer addition time was determined by online monitoring of medium glucose consumption. Therefore, beginning after 24 hours of further cultivation 20 µL medium samples were taken every 12 hours, quenched with 30 % ice cold acetonitrile and measured using hydrophilic interaction chromatography (HILIC).

For HILIC measurements a HILIC SeQuant® ZIC®-pHILIC column (100 x 2.1 mm i.d., 5 µm particle size, Merk Millipore, Burlington, MA, USA) was used. Column oven temperature was set to 35 °C and a sample injection volume of 1 µL was set. Liquid phase was applied at a constant flow rate of 300 $\mu L \cdot min^{-1}$ and consisted of 10 mM ammonium formate with pH 6 (eluent A) and acetonitrile with 5 % ammonium formate ($v \cdot v^{-1}$) (eluent B). A gradient elution was applied starting with 100 % B for 2 min with a subsequent 14 min linear decrease to 58 % B followed by constant 58 % B for 2 min and finally a one min linear gradient to 100 % B and 100 % B for one min.

The tracer was then added along with additional fresh medium after an exponential increase in glucose consumption was observed. Further 20 µL samples were taken every 12 hours for online monitoring of glucose and tracer concentrations in the culture medium. Cultivation was stopped after glucose was depleted.

After the cultivation was stopped, two different types of samples were prepared. On the one hand, culture supernatant samples primarily reflecting the exometabolome (extracellular metabolites) and mycelium extracts that reflect rather the endometabolome (intracellular metabolites).

### 1.2 Fungal strains and cultivation conditions

*Fusarium graminearum* PH-I was grown on liquid minimal medium (1.0 g·L$^{-1}$ $KH_2HPO_4$, 0.5 g·L$^{-1}$ $MgSO_4 \cdot 7 H_2O$, 0.5 g·L$^{-1}$ KCl, 0.48 g·L$^{-1}$ $NH_4NO_3$, 0.2 mL·L$^{-1}$ Vogel's trace solution [1]) containing 1 % (w·v$^{-1}$) D-glucose (native glucose for the standard tracer approach and U-$^{13}C_6$ labeled D-glucose for the reversed tracer approach) on 24 well cell culture plates in a volume of 1.0 mL per well. In each case 1 mL medium was inoculated with 6 µL fresh spore solution (final concentration 25,000 spores·mL$^{-1}$). After 72 hours pre-cultivation the tracer solution, consisting of the respective minimal medium containing 0.5 % (w·v$^{-1}$) of the corresponding tracer, was added to the cultures in a volume of 1.0 mL per well. Cultivation was then continued under identical conditions. The cultivation was stopped after 96 hours, where remaining glucose concentration reached 0.1 % (w·v$^{-1}$).

*Trichoderma reesei* strains QM6aΔ*tmus53*, QM6a Δ*tmus53*Δ*sor1* and QM6a Δ*tmus53ReYpr1* were grown from spore solutions, stored on -80 °C, on 2 % (w·v$^{-1}$) malt extract (MEX) agar with 0.5 % (w·v$^{-1}$) peptone at 30 °C and 80 % relative humidity. Fresh spores were scraped off the plates and suspended in $ddH_2O$ and filtered through glass wool. The spore solution obtained were adjusted with $ddH_2O$ to an $OD_{700}$ of 0.5 absorption units at 1 cm path length. A minimal medium based on Mandels-Andreotti (MA) medium [2] without peptone containing 1 % (w·v$^{-1}$) D-glucose (native glucose for the $^{13}C$-labeled tracer and U-$^{13}C_6$ labeled glucose for the native tracer) was then used for pre-cultivation of *T. reesei* on 24 well cell culture plates in a volume of 1.0 mL per well, inoculated with 100 µL spore solution at 30 °C, 80 % relative humidity in darkness for 48 hours. Six biological replicates were made for each approach. After this pre-cultivation the tracer solution, consisting of the respective MA medium containing 0.5 % (w·v$^{-1}$) of the corresponding tracer, was added to the cultures in a volume of 1.0 mL per well. Cultivation was then continued under identical conditions. The cultivation was stopped after 72 hours, where the concentration of remaining glucose reached 0.1 % (w·v$^{-1}$).

### 1.3 Sample preparation

Culture medium supernatant samples and fungal mycelium extract samples. First the mycelium was removed and transferred in 2 mL tubes. The supernatant was filtered through glass wool to remove mycelium left overs. The filtered supernatant was cooled on ice and quenched by adding 30 % (v·v$^{-1}$) cold (-20 °C) acetonitrile, centrifuged at 30,000 x g for 20 minutes at 4 °C and transferred into HPLC vials. The mycelium was washed twice with 1.5 mL $ddH_2O$ each. After each washing step, the mycelium was carefully centrifuged at 1,500 x g for 5 minutes at 4 °C and the washing water was discarded. Then two steel balls (2 mm diameter) were added to each tube as well as 600 µL ice cooled extraction solvent consisting of methanol and $ddH_2O$ in a ratio of 1:1 (v·v$^{-1}$). The mycelium was then ground using a Retch mill at 30 Hz in pre-cooled (-20 °C) tube holder blocks for 5 minutes. Then 400 µL $ddH_2O$ were added per tube to achieve a proportion of 30 % organic solvent in the extracts. The samples were vortexed roughly and finally centrifuged at 30,000 x g for 20 minutes at 4 °C and the supernatant was transferred into HPLC vials.

For the stable isotope assisted global labeling approach for the detection of all features corresponding to ions of metabolites, actually originating from the respective fungus by means of MetExtract II [3] data evaluation, corresponding samples of fungi grown on native glucose and in parallel grown on U-$^{13}C_6$ labeled glucose without the addition of a labeled tracer were mixed in each case in a ratio of 1:1 as described in Bueschl et al. [4].

### 1.4 LC-HRMS measurement

A Vanquish Duo UHPLC system (Thermo Fisher Scientific, San Jose, CA, USA) with both an reversed-phase (RP) XBridge BEH $C_{18}$ column (150 × 2.1 mm i.d., 3.5 µm particle size, Waters, Milford, MA, USA) coupled with a pre-column ($C_{18}$ 4 × 3 mm i.d., Security Guard Cartridge, Phenomenex, Torrance, CA, USA) coupled to an Orbitrap QExactive HF (Thermo Fisher Scientific) equipped with an electrospray ionization source (HESI) was used for separations and data acquisition.

For measurements the MeOH/$H_2O$ HPLC method as described in Kluger et al. [5] was used. Only the sample injection volume was 5 µL instead of 10 µL.

The HESI ion source was operated in fast polarity switching mode using following MS and HESI parameters: Scan range: *m/z* 100-1500; Sheath gas flow rate 55 arb. units; Aux gas flow rate 5 arb. units with a temperature of 350 °C, spray voltage 3.5 kV (positive mode) or 3.0 kV (negative mode); Full MS mode, R=120.000, AGC target 3E6.

### 2 Supplementary figures

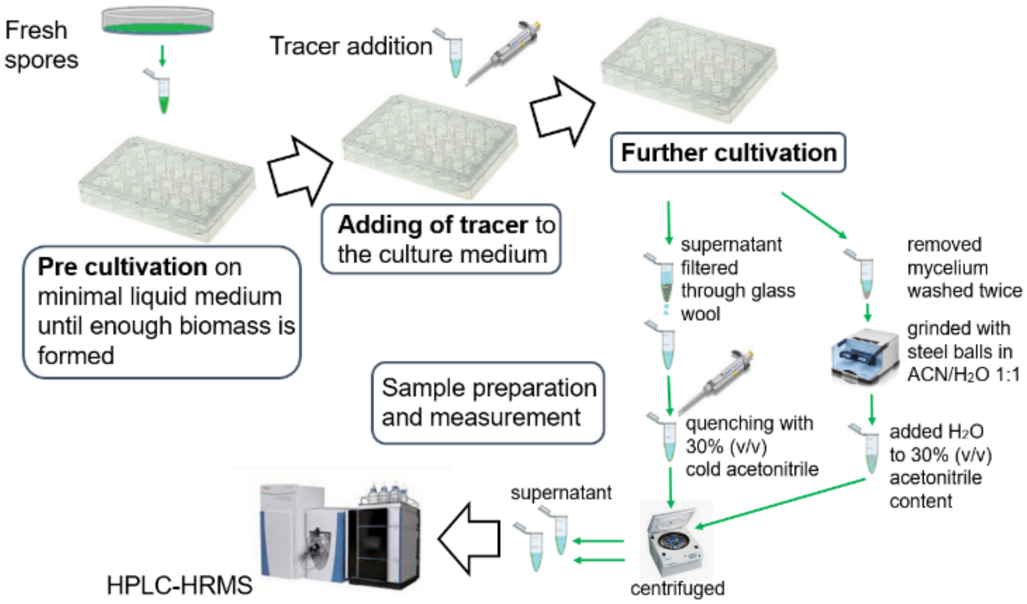

**Figure S1.** Schematic representation of the SIA tracer approach workflow used to obtain the biological samples which led to the data sets used in the presented proof of concept examples for the applicability of the CPExtract algorithm.

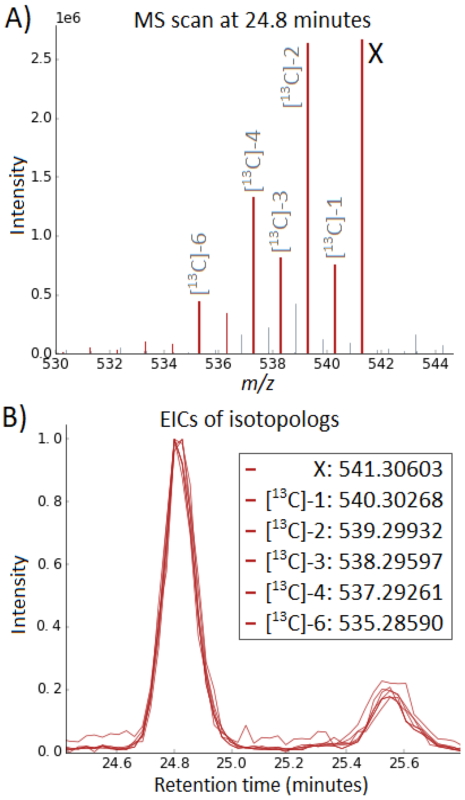

**Figure S2**. Sample mass spectrum and EICs of a tracer derived isotopolog pattern that was searched for with CPExtract. "X" represents the uniformly $^{13}$C-labeled metabolite ion (i.e. all carbon atoms are $^{13}$C), while the [$^{13}$C]-n represent the exchange of n $^{13}$C isotopes with n $^{12}$C isotopes. A) Mass spectrum showing the expected isotopolog pattern of a polyketide with at least three $^{12}$C-malonate units. B) The different isotopologs illustrated in A show perfect co-elution when their EICs are overlaid.

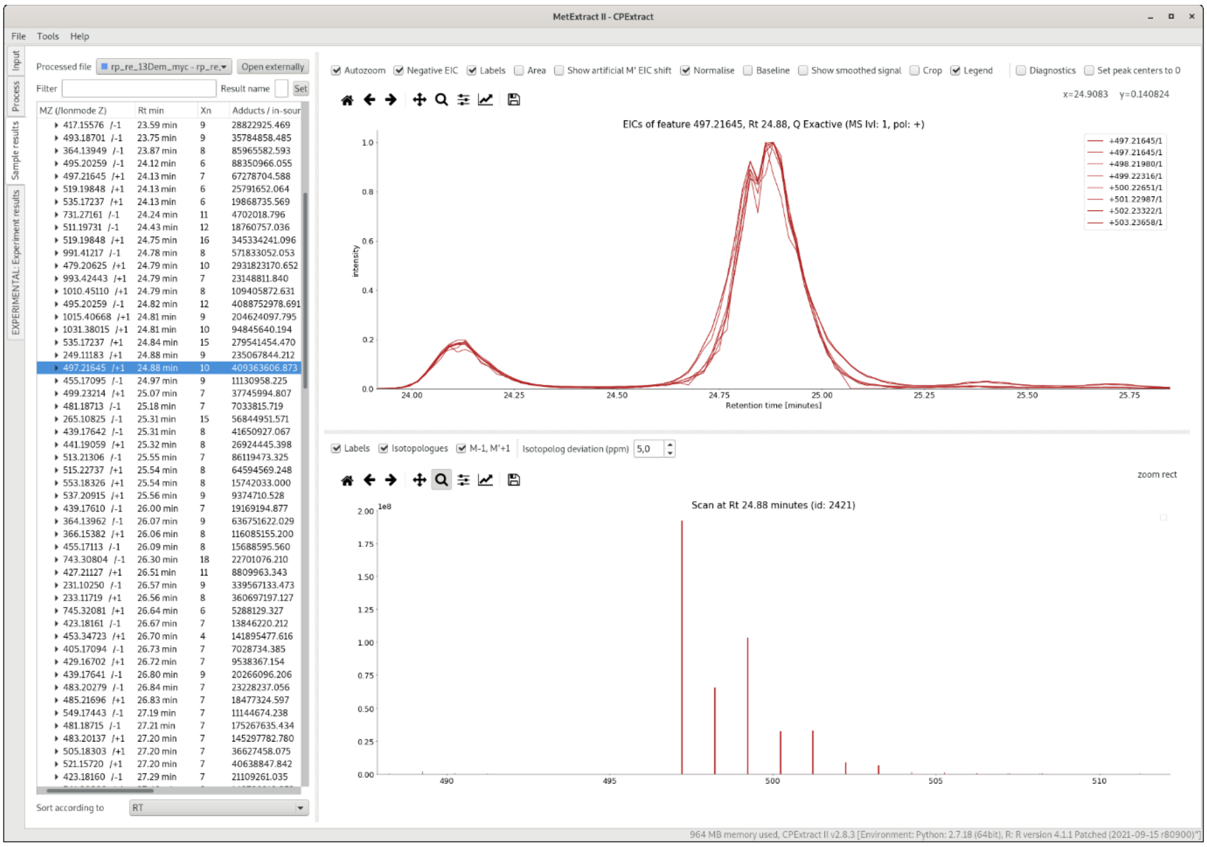

**Figure S3.** Screenshot of CPExtract software results panel showing an example of a detected tracer derived metabolite ion and its respective isotopologs in form of overlaid, normalized EICs (right side, upper plot) and the spectrum with rule-compliant isotopologs marked in red (right side, lower plot).
